## Supplemental Text for "Dynamic collateral sensitivity profiles highlight challenges and opportunities for optimizing antibiotic sequences"

### Supplemental Material: Temporally dynamic collateral sensitivity profiles complicate optimal dosing strategies

The supplemental material contains Supplemental Text, 6 supplemental figures, and Supplemental Tables (xlsx file).

#### Multi-drug sequences can minimize cumulative resistance to the applied drugs

While our experiments are limited to a single drug switching event, we also wondered whether these measurements might provide a null model for more complex treatment strategies that involve multiple drugs (Fig S6). Intuitively, the model corresponds to the accumulation of multiple mutations whose effects combine approximately independently, as described below. We stress, however, that this is a strong assumption, and other evolutionary processes are certainly possible (if not likely), so this should be viewed, at best, as a null model that essentially describes the combinatorial patterns associated with the measured collateral profiles.

We consider a simple model where collateral effects accumulate linearly based on the resistance values measured for the selecting drug in question [40]. Because collateral effects are measured on a log scale, this linear model reduces to one where fractional changes in resistance levels are multiplicative [75]. More specifically, in a system with $N$ drugs, the state at time $t$ is defined by an $N-$dimensional vector $S_t$, with each component of that vector the collateral value (i.e. log-scaled fold change in $\mathrm{IC}_{text50}$) associated with a specific testing drug. The state of the system at the next time point, $t+1$, is then given by $S_{t+1} = S_t + \delta C$, where $\delta C$ accounts for the change in collateral profile due to selection in the currently applied drug. When an antibiotic is used for the first time, $\delta C$ is randomly drawn from one of the 4 (replicate) profiles measured on day 2 (i.e. one column of the matrix in Fig. 2B, upper left). However, for each subsequent use of that antibiotic, $\delta C$ is given by the collateral profile corresponding to the next temporal step on the chosen trajectory (i.e. by the *same* column but in the measured collateral matrix at the next time step). For example, the first time a population is exposed to CIP, $\delta C$ is drawn randomly from one of the 4 collateral profiles we observe on day 2 of CIP selection (for example, column 3 in Fig 2B, upper left matrix). However, when the population is exposed to CIP again, $\delta C$ will be given by the same column (column 3) of the collateral matrix measured on day 4 (Fig 2B, upper right). The goal of this model is to capture the *measured* temporal changes associated with repeated exposures to the same drug while neglecting higher-order temporal effects between different drugs applied multiple times (as measuring these higher-order effects would require an exponentially growing number of measurements because of the combinatorial explosion in possible trajectories). While this model is clearly an oversimplification of the full evolutionary dynamics, we have shown similar models can offer qualitatively accurate predictions in relatively short evolution experiments [40,44]. And because of the model's simplicity, we can exhaustively simulate all possible trajectories in hopes of identifying candidate drugs for further experiments.

Using this model, we simulated 8-day trajectories using every possible combination of 5 drugs, with drugs potentially alternated every 2 days (days 0, 2, 4, and 6). Using these results, we calculated the "applied drug resistance", which we define as a cumulative measure of the collateral effects for the applied drugs across all time-steps. It is given by $R_{\mathrm{app}} = \sum_n S_{j_n}(n)$, where the sum runs over all time steps of the treatment and $S_{j_n}(n)$ is the component of the state vector corresponding to the applied drug $j_n$ at each step (Fig S6A). Large positive values of $R_{\mathrm{app}}$ correspond to treatments with, on average, high levels of resistance to the applied drugs, while large negative values correspond to high levels of sensitivity.

We find that the applied drug resistance $R_{\text{app}}$ depends on both a) the number of different drugs in the sequence and b) the specific, time-dependent ordering of those drugs (Fig. 3A). As the number of drugs used in the policy increases from 2 to 4 (Fig. 3A top to bottom, black curves), the distribution of $R_{\text{app}}$ narrows and the mean decreases. Intuitively, the decreasing mean is perhaps not surprising, as switching between multiple drugs is expected to create an increasingly difficult evolutionary challenge. Surprisingly, however, at a global level both the best (smallest $R_{\text{app}}$) and worst (largest $R_{\text{app}}$) schedules require switching between multiple drugs, indicating that multiple drug sequences (black) are not always an improvement over single (constant) drug schedules (red curves).

#### Switching drugs may raise global resistance to the set of available drugs.

We next used simulation data to calculate the "total resistance", which we define as the cumulative sum of all entries of the state vector over the treatment period, $R_{\text{tot}} = \sum_n \sum_j S_j(n)$, where $S_j(n)$ is the $j$-th entry of the state vector at time point $n$. This quantity is a global measure of resistance or sensitivity to all available drugs. In contrast to $R_{\text{app}}$, the schedules that lead to the lowest levels of $R_{\text{tot}}$ (total resistance) all correspond to single-drug schedules (Fig. 3B). Despite the fact that single drug schedules are unable to exploit collateral sensitivities that may arise, they are more effective at minimizing global resistance to the pool of drugs, in part because early stages of adaptation are dominated by costly collateral resistance.

#### Drug timing frequently impacts applied drug resistance but not total resistance

To evaluate the impact of the timing and number of drug switches in a treatment protocol, we compared the best (smallest value of $R_{\text{app}}$ or $R_{\text{tot}}$) and worst (largest value of $R_{\text{app}}$ or $R_{\text{tot}}$) treatment outcomes for an exhaustive combination of every 2-drug treatment in our study (Fig S6C-D). We found that even when restricted to policies involving only 2 drugs, the best schedule can dramatically outperform the worst schedule in terms of applied drug resistance, suggesting that the timing of the switches, not merely the drugs used, can have a significant impact. For example, for the drug pair DOX-CRO, using the pair optimally leads to a small applied drug resistance score ($R_{\text{app}} \approx 3.6$), whereas using the drug pair sub-optimally leads to a significantly larger score ($R_{\text{app}} \approx 14$). On the other hand, when it comes to the total collateral profile resistance (Fig S6D), the timing of the drug switches has a relatively small impact for all drug pairs.

It is important to remember that we used an extremely simple additive model of resistance to simulate possible evolutionary trajectories. While additive models can be useful to guiding experiments, particularly on short timescales [40,44], and have shown to be surprisingly robust in some contexts [76], we expect them to ultimately fail when epistasis or evolutionary hystersis are strong [77]. We therefore reiterate that the primary purpose of such simple models is to generate hypotheses to guide experiments, not to provide a true theoretical description of the system.

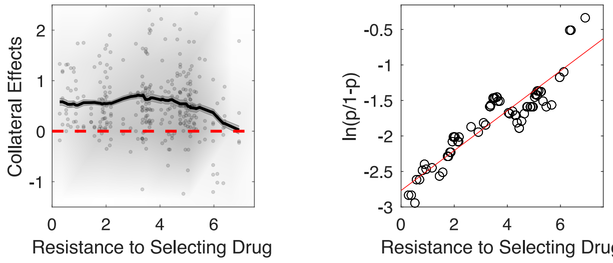


**Fig S1. Cumulative collateral effects exhibit early resistance but trend toward sensitivity with adaptation.** Left panel: cumulative collateral effects (relative to ancestor strain) as a function of resistance to selecting drug. Dots are individual populations, shading indicates relative density of points, and curve is moving average (shading is ± standard error over each window). Right panel: probability of instantaneous sensitivity ($p$) varies with cumulative resistance to selecting drug. Line is fit from logistic regression (i.e. fit of $\ln(p/(1-p)) = c_1 + c_2 R$, where $R$ is resistance to selecting drug and $c_1$ and $c_2$ are intercept and slope parameters, respectively). Slope parameter: $c_2 = 0.3$ ($(0.08, 0.50)$, 95 percent confidence interval); $p_{\text{value}} = 0.006$. Averages and probabilities are calculated over sliding windows of size 2 (in units of resistance to selecting drug).

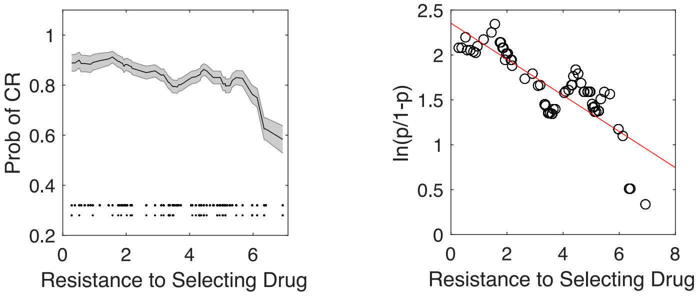


**Fig S2. Collateral resistance decreases as resistance to selecting drug increases.** Left panel: probability of (instantaneous) collateral resistance (CR) as a function of resistance to selecting drug. Curve is a moving average (shading is ± standard error over each window). Lower inset points show individual data points (top row is CR, bottom row is not CR) used to calculate probability. Right panel: probability of resistance ($p$) varies with resistance to selecting drug. Line is fit from logistic regression (i.e. fit of $\ln(p/(1-p)) = c_1 + c_2 R$, where $R$ is resistance to selecting drug and $c_1$ and $c_2$ are intercept and slope parameters, respectively). Slope parameter: $c_2 = -0.2$ ($(-0.4, -0.01)$, 95 percent confidence interval); $p_{\text{value}} = 0.04$. Averages and probabilities are calculated over sliding windows of size 2 (in units of resistance to selecting drug).

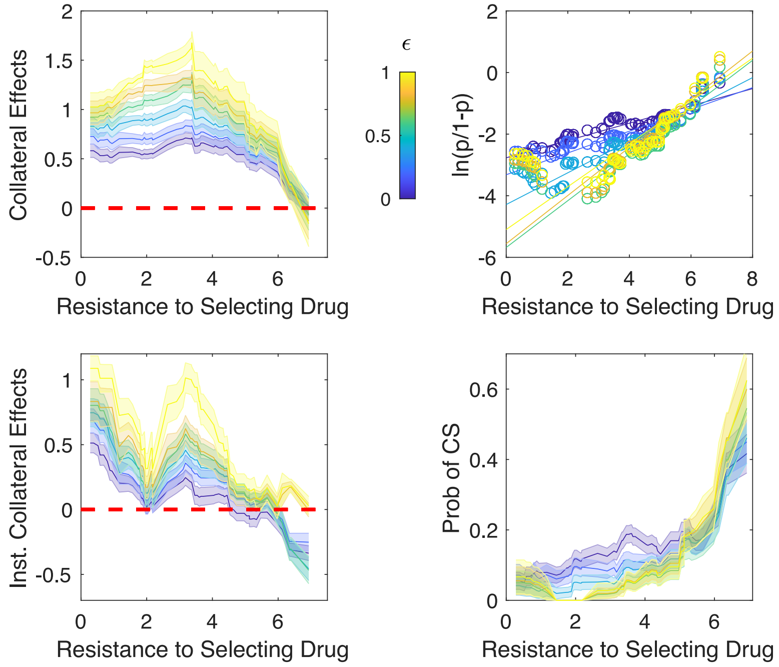


**Fig S3. Modulating threshold value does not change qualitative features of collateral effects** Collateral effects (top left), logit function (top right), instantaneous collateral effects (bottom left), and probability of collateral sensitivity (bottom right) as a function of resistance to selecting drug. All collateral effects (i.e. log of fold change in $IC_{50}$) with an absolute value less than $\epsilon$ are removed prior to analysis. Different colors represent the same analysis but with different values of $\epsilon$. All moving averages are taken over a window size of 2. Compare to Figures 2, S1. For logistic regressions, $p_{\text{value}} < 0.05$ with a positive slope parameter (i.e. frequency of CS increases) for all values of $\epsilon$.

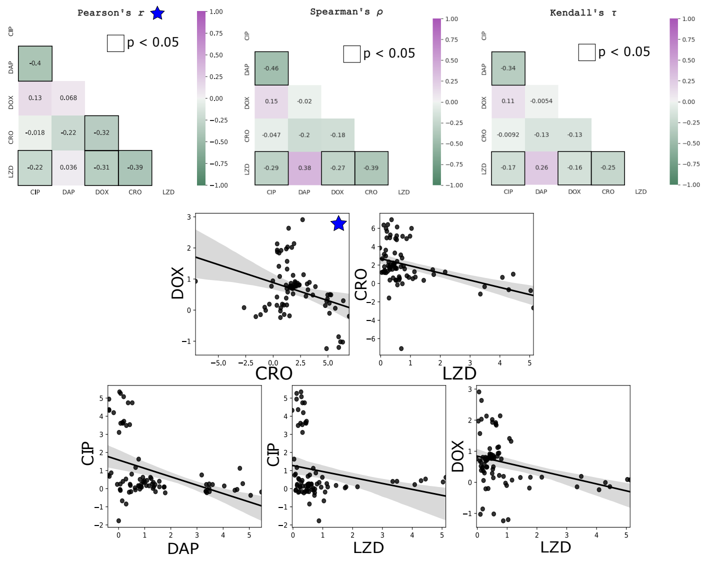


**Fig S4. Correlation between collateral effects between different testing antibiotics. Top panels:** Pearson, Spearman, and Kendall correlation coefficients between collateral profiles between each of the 5 testing antibiotics. Dark squares highlight correlations with statistically significant correlations ($P < 0.05$), whose distributions are shown as scatter plots. **Bottom panels:** pairwise scatter plots of resistance profiles selected by different testing antibiotics; only pairs with significant correlations ($P < 0.05$) are shown. Each point is the measured resistance to each of the single antibiotics labeled on the axes. DOX-CRO pair (blue star) is only statistically significant using Pearson's correlation.

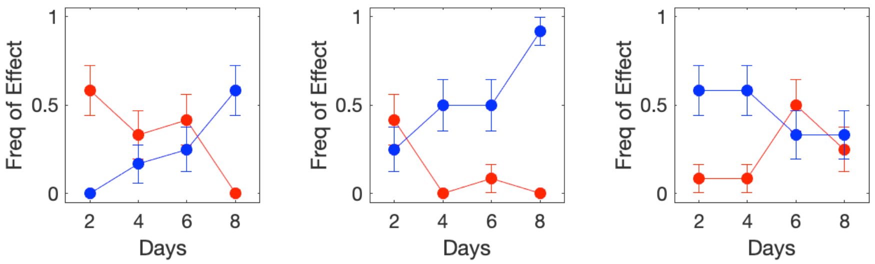


**Fig S5. Frequency trends within individual populations do not depend sensitively on threshold criteria for defining collateral effects changes.** Frequency of sensitivity (blue) or resistance (red) for different time points from populations adapted to LZD (left), CRO (middle), and CIP (right) and exposed to CRO (left), DOX (middle), and CRO (right). In this plot, collateral effects are considered significant if errorbars ($\pm$ 1 standard error) do not overlap. See also Figure 3 C.

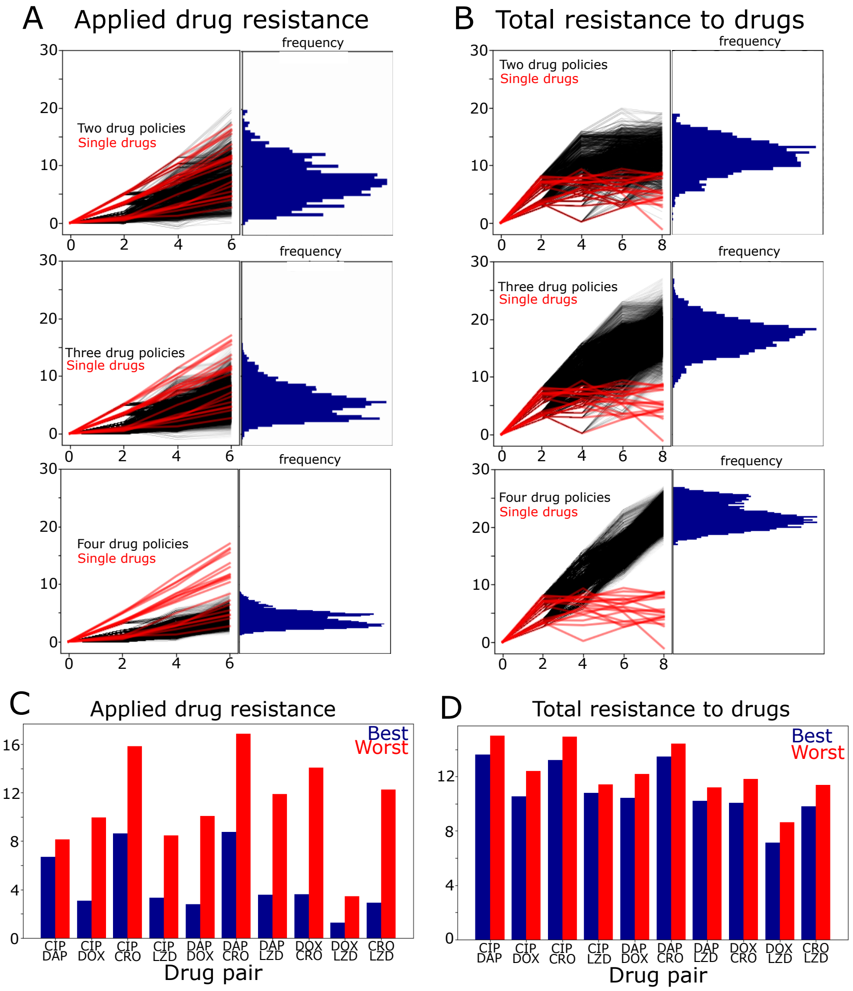


**Fig S6. Simulations suggest optimal dosing can be aided or hindered by temporally resolved collateral profiles.** **(A)** The left panel in each figure shows simulated resistance level to the applied drug (vertical axis) over time (horizontal); the right panel shows a histogram of frequencies across all trajectories. Plots include exhaustive simulations of every hypothetical trajectory assuming any of the five drugs can be chosen at days 0, 2, 4 and 6. If the population is exposed to the drug for the first time, any of the four measured replicates are accessible to the population. However, once a population has chosen a trajectory in an antibiotic, further use of that antibiotic will stay on that chosen trajectory. For example, a dose schedule of CIP-CIP-CIP-CIP would result in selecting one of the four CIP replicates measured in the heatmap corresponding to days 0-2 evolution (e.g. CIP2), followed by that replicates measured results for days 2-4, 4-6 and 6-8. A running total of the resistance of the population to the applied drug is plotted both for the trajectories where more than one drug is chosen (black) and for the single drug trajectories (red). The top box plots all two drug policies, the middle box plots all three drug policies and the final box plots all four drug policies. **(B)** Same as panel A except vertical axis is the global resistance (the sum of the collateral profile across all drugs). **(C)** The best and worst performing drug-cycling policy for each drug pair is shown using the running total of resistance to the applied drug (as in A). **(D)** The best and worst performing drug-cycling policy for each drug pair is shown using the sum collateral profile as the metric (as in B).
